# Estimating stem cell fractions in hierarchically organized tumors

**DOI:** 10.1101/013672

**Authors:** Benjamin Werner, Jacob G. Scott, Andrea Sottoriva, Alexander R.A. Anderson, Arne Traulsen, Philipp M. Altrock

## Abstract

Cancers arise as a result of genetic and epigenetic alterations. These accumulate in cells during the processes of tissue development, homeostasis and repair. Many tumor types are hierarchically organized and driven by a sub-population of cells often called cancer stem cells. Cancer stem cells are uniquely capable of recapitulating the tumor and can be highly resistant to radio-and chemotherapy treatment. We investigate tumor growth patterns from a theoretical standpoint and show how significant changes in pre-and post-therapy tumor dynamics are tied to the dynamics of cancer stem cells. We identify two characteristic growth regimes of a tumor population that can be leveraged to estimate cancer stem cell fractions *in vivo* using simple linear regression. Our method is a mathematically exact result, parameter free and does not require any microscopic knowledge of the tumor properties. A more accurate quantification of the direct link between the sub-population driving tumor growth and treatment response promises new ways to individualize treatment strategies.

**Significance Statement:** Under the cancer stem cell hypothesis a tumor population is driven by a fraction of self-renewing cancer stem cells. Absolute and relative size of this population in human cancers at any stage of the disease remains unknown. We formulate a mathematical model that describes the tumor cell population’s growth dynamics and response to therapy. This allows to estimate cancer stem cell fraction from longitudinal measurements of tumor size (often available from imaging). Such estimates are critical because treatment outcome and risk of relapse depend on the tumor’s capacity to self-renew. Ideally, by tailoring patient treatment strategies based on the relative abundance of cancer stem cells could lead to radically different therapeutic regime and to the successful eradication of the disease.

## 1 Introduction

Cancer comprises a group of diseases that involve abnormal and uncontrolled proliferation of cells that were once normal. These aberrant properties are induced by alterations in genes that control cell regulatory mechanisms, microenvironmental response and cell-cell signaling: a group of functions referred to as the ‘Hallmarks of Cancer’ [1]. Although large-scale genomic studies have revealed the spectrum of genomic profiles in many cancers [2], recent accumulating evidence shows that cancers are characterised by extensive inter-patient [3] and intra-tumor heterogeneity [4] as a consequence of tumor evolution [5]. This heterogeneity bridges multiple scales and is not only tied to the tumor but also the context within which it grows, its microenvironment [6].

In addition, tumors are often comprised of cancerous cells in distinct stages of differentiation [7]. This “phenotypic” diversity likely is a remainder of the hierarchical organization of the tumors’ tissue of origin. In most healthy tissue, stem cells maintain tissue homeostasis and a certain number of cell differentiation compartments give rise to the production of phenotypically distinct mature cell types [8, 9]. The finding that tissue organization can be maintained in tumors has lead to the postulation of the existence of cancer stem cells, termed the cancer stem cell hypothesis. Under this hypothesis a fraction of cells are uniquely able to seed, maintain and re-seed tumors [7]. First identified in leukemia [8], cancer stem cells have since been shown to drive a number of solid tumors, including colon [10, 11, 12], brain [13], breast [14], head and neck [15], lung [16], and melanoma [17], among others.

These *in vivo* and *in vitro* observations are complimented by a rich emerging body of literature using a variety of mathematical methods to model the hierarchical organization of tissues in physiological and pathological contexts. Tissue-specific models include those focused on the cell hierarchy in colonic crypts [18, 19, 20] and leukemias [21, 22]. Other studies have sought to understand the general dynamical behavior of tissues organized in a hierarchical way [23, 24, 25, 26].

Finding an effective treatment strategy against cancers organized into a hierarchy is thought to require the elimination of all cancer stem cells [27, 28]. However, an increase of the cancer stem cell fraction during treatment is frequently observed [29, 30]. This increase is potentially due to various mechanisms, such as cancer stem cell quiescence [31], specific intrinsic mechanisms of radioresistance[32] and chemoresistance [33], or even microenvironmental plasticity of the non-stem phenotype [34, 35] promoting the cancer stem cell population.

Naïvely, the specific targeting of cancer stem cells seems a promising and necessary approach to improve treatment [5]. Different sizes of cancer stem cell populations between patients could potentially influence individualized treatment strategies and improve prognosis [36]. Unfortunately, the fraction of tumor-driving stem cells is unknown at diagnosis. Currently, the only method that exists that can infer this information for specific patients is from direct biopsy before and after treatment. This has major limitations due to marker resolution and sampling frequency and is further confounded by location dependence and intrinsic heterogeneity, not to mention risk to the patient. Ideally, we require a continuous measure of the stem fraction that can easily be obtained from relatively non-invasive means.

We use a multi-compartment approach, which allows an analytical description of cell population dynamics in hierarchically organized tumors [37, 38]. It had been noted before that hierarchical tumors transition from a fast into a relatively slower phase of tumor growth. During treatment response, particularly in targeted treatment of leukemias, a similar effect of strong response followed by weaker response is common [39, 40, 41]. Here we show that these transitions are tied to the dynamic characteristics of cancer stem cells. Moreover, our analytical results allow us to exploit this universal property of hierarchical tumor organization. We show how one can estimate the fraction of tumor-driving cancer stem cells from purely macroscopic observables, such as information about tumor size gleaned from medical imaging.

## 2 Results

We model hierarchical tumor organization by a multi compartment approach (see Fig. 1). Each compartment represents cells at certain differentiation or proliferation stages [21, 37]. We investigate a minimal model, where compartment 0 contains stem cells that proliferate at a rate *χ*_S_ and die at a rate *d*_S_. Transient amplifying cells proliferate at a rate *χ*_D_ and die at rate *d*_D_. Self-renewal of cancer stem cells occurs with a probability *p*. Differentiation into transient amplifying cells occurs with probability 1 – *p*. The cell lineage can undergo at most *m* cell doublings before the most differentiated cells enter senescence. This resembles a cells’ Hayflick limit, which might be a consequence of critically short telomeres, or other cell regulatory mechanisms [42]. Usually, the proliferation rate of transient amplifying cells is increased as compared to stem cells, for example we have *χ*_S_ < *χ*_D_. Modifying the death rates allows us to implement a minimal representation of immune-response during tumor growth, as well as different treatment regimens.

**Figure 1.**
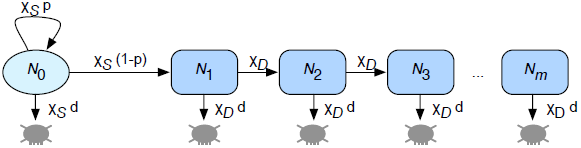
Model schematic, showing key parameters governing the mathematical model based on the cancer stem cell hypothesis. Cancer stem cells, denoted by *N*_0_, are exclusively able to maintain the tumor cell population. Transient amplifying cells (*N*_1_,…, *N*_*m*_) undergo *m* cell division before which they enter cell senescence. Cancer stem cells proliferate with a rate *χ*_S_, self renew with probability *p* and die at a rate csd. Transient amplifying cells proliferate with rate *χ*_D_ and die at a rate *χ*_D_*d*.

The deterministic dynamics of a cell population that is organized in such a hierarchy can be described by a set of coupled linear differential equations. The general analytical solution reduces to a set of weighted exponential functions (see Methods for details). If there are a finite number of cell divisions, the tumor growth curve decomposes into two regimes. The first regime is driven by cancer cells filling up compartments of higher differentiation. The second phase is characterized by a dynamic equilibrium, in which this drive from below is balanced by loss of cells due to senescence. In addition, and more importantly, we can infer the impact of the fraction of cancer stem cells on possible ‘phase transitions’ during tumor growth and response to treatment.

### Tumor growth

We initiate tumor growth with a single (cancer stem) cell in compartment 0. This is in line with the cancer stem cell hypothesis [43] and does not necessarily imply that the cell of origin was a stem cell. Potentially stem like properties can be acquired at later stages of the hierarchy, as is common in for example different acute leukemias [8, 28]. The proliferation parameters of cancer stem cells determine the long term behavior of the tumor. A tumor grows continuously (and potentially becomes a detectable cancer), if *p* > (1 + *d*)/2. The probability of stem cell self renewal *p* needs to be sufficiently large to compensate loss of stem cells by random cell death *d*. In contrast, the tumor population vanishes for insufficient cancer stem cell self renewal. In the following, we assume that *p* is sufficiently large to allow for a growing tumor.

In this scenario, the population of cancer stem cells expands. In addition, cancer stem cells differentiate into transient amplifying cells that comprise the bulk of the tumor. However, after sufficient time, loss of differentiated cells due to cell death or cell senescence and gain of differentiated cells due to doublings of transient amplifying cells balance one another. When this balance is reached, the tumor growth dynamics transition into a second phase. Further tumor growth is limited by the expansion rate of the cancer stem cell compartment, see Fig 2 for an example. These two distinct growth phases are a generic property of hierarchically organized tumors. WE only assume that proliferation rates of tumor initiating cells and further differentiated cells differ [38].

**Figure 2.**
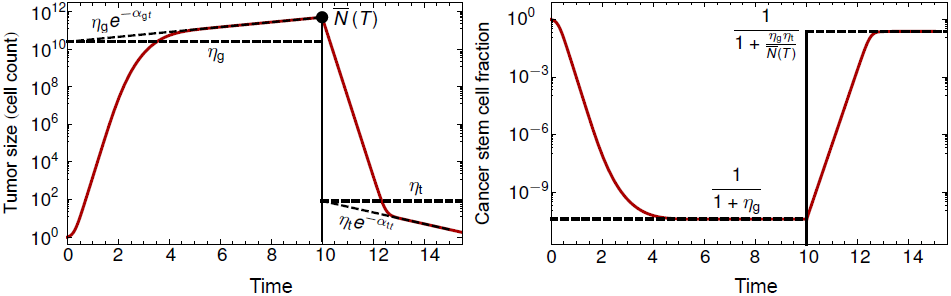
Inferring the fraction of cancer stem cells from tumor growth curve or treatment response. The estimation method is an analytical result that follows from our mathematical model. The red line shows one realization of the model for tumor growth and treatment response. The dashed lines correspond to exponential fits and their offsets that do not require any detailed knowledge on the tumor cell properties. If the offsets *η*_*g*_ and *h*_*t*_ can be estimated from the regression during growth (*g*) and treatment (t) respectively and the tumor size at the beginning of treatment is *N*̅, then one can infer the equilibrium fraction of cancer stem cells during tumor growth and treatment response from purely macroscopic observables without the need of detailed knowledge of tumor cell properties.

### Cancer stem cell fraction

The fraction of cancer stem cells *r*_*g*_ at time *t* during tumor growth is given by

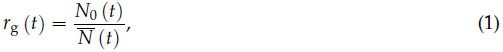

where *N*_0_(*t*) corresponds to the number of stem cells, and is *N*̅(*t*) the sum of all tumor cells at time *t*. The fraction of tumor stem cells decreases in the first phase of tumor growth. Then it evolves towards an equilibrium state for any possible combination of cell proliferation parameters, see Fig. 2.

The value of the cancer stem cell fraction in dynamic equilibrium depends on the sign of the differential flow between the stem and the non-stem compartments. This flow describes the difference between the net expected growth of the cancer stem cell compartment and the net loss in any given differentiated (non-stem) compartment due to cell differentiation or death.

If the differential flow is negative, more stem cells are lost by differentiation than gained by self renewal. Then the stem cell fraction tends to zero. If the differential flow is exactly zero, cancer stem cells furnish half of the tumor population. For positive values of the differential flow, we have a surplus in the production of differentiated tumor cells as compared to their losses during further differentiation. In this third case, one can show that the time dependent components converge to a constant value (*N*̅(*t*)/*N*_0_(*t*) → const.) and the fraction of cancer stem cells take a non trivial value between 0 and 1, see Methods for details. Thus, the relative composition of the growing tumor remains constant in the second phase of tumor growth and is given by

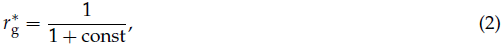

see Fig. 2 for an example. The value of the constant is calculated analytically in the Method section and involves all model parameters. Thus from the model’s perspective, a detailed microscopic knowledge of a tumor’s properties seems to be a prerequisite to estimate 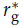. For example, all proliferation parameters of different cancer cell types should play a role. Yet, this detailed knowledge is unlikely or even impossible to gain in a clinical setting. In the following we propose an alternative method to estimate the constant in the denominator of Equation (2).

### Estimating the cancer stem cell fraction during growth

An important macroscopic observable of a tumor is its growth curve. Small changes in tumor sizes can be visualized and analyzed effectively, for example by high resolution magnetic resonance imaging. Thus, tumor growth curves can be assessed reliably within relatively short time intervals, and are recorded routinely in modern clinical care. Most importantly, these techniques do not require any knowledge of the microscopic tumor properties.

Our mathematical model provides an analytical description of the tumor growth curves. The first phase of tumor growth is a combination of rapid stem and differentiated cell proliferation. However, as the tumor reaches an equilibrium state its growth follows a slower exponential expansion rate driven by the stem cell pool. In dynamic equilibrium, tumor growth can be captured analytically by a single exponential function of the form *a* e^*bt*^. The coefficients *a* and *b* involve the parameters of the mathematical model.

Interestingly, one finds that the coefficient *a* coincides with the constant in the exact expression of 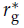 and we can write 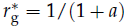. This observation allows us to estimate 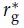 from tumor growth curves. Instead of calculating the parameter *a* analytically, one can fit an exponential function *η*_g_e^*α*_g_*t*^ to a growth curve of a tumor in equilibrium via linear regression of the logarithmically transformed tumor size data. This fit gives two parameters *α*_g_ and *η*_g_, which do not require any detailed microscopic knowledge. Moreover, the offset *η*_g_ of the regression corresponds to the theoretically calculated parameter *a*. This observation allows us to estimate the fraction of tumor driving cancer stem cells via the relation

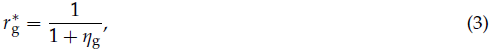

from an exponential fit *η*_g_e^*α*_g_*t*^ to the tumor growth curve, see Fig. 2 for an illustrative example.

### Estimating the cancer stem cell fraction under treatment

Our general approach allows us to implement treatment strategies by altering the death rates *d* (or other parameters of the model) of cancer cells. A hierarchically organized tumor shrinks continuously under treatment if the death rate d of cancer cells exceeds the self renewal capability *p* of stem cells by *d* > 2*p* – 1, and no treatment resistant clone is present [44]. Under continuous treatment, like unperturbed growth, we observe a bi-phasic response. The tumor cell population shrinks fast initially, and transitions into a slower decrease after a characteristic time.

The first phase is dominated by the death of differentiated cells. In this phase, treatment selects for cancer stem cells, see Fig. 2. Then the tumor reaches a dynamic equilibrium stage, in which the relative flux of cell renewal and cell loss balance. The relative composition of the tumor remains constant, despite a continuous decrease in tumor size. This causes the transition into a second phase of tumor shrinking, where the initial treatment effect starts to diminish.

The stem cell fraction that is active during treatment response can be estimated by an exponential fit *η*_t_e^*α*_t_*t*^ to the tumor shrinkage curve. Under treatment, the tumor’s initial condition is not a single seeding cancer stem cell. We have to take all cancer cells into account. This changes our estimates and introduces additional complexity. In the Methods section we show that *η*_t_, the offset of the exponential fit under treatment, *η*_g_, the offset of the exponential fit during growth and *N*̅(*T*), the total tumor size at treatment initiation *T* suffice to accurately estimate the fraction of cancer stem cells under treatment, which is then given by the relation

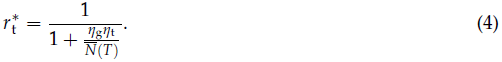

This is an exact result of our model and its structure is very similar to the case of untreated tumor growth, see Fig. 2. However, we require additional information to estimate the fraction of cancer stem cells under treatment, but this information can be gained from macroscopic observations with no need for detailed knowledge about the microscopic tumor properties.

## 3 Discussion

The cancer stem cell hypothesis was formulated almost two decades ago and has attracted much attention and research but also many critics and much skepticism ever since [45]. While the existence of cancer stem cells in some tumors is well established, the situation in other cancers remains somewhat unclear [9, 43]. However, its impact on the understanding and treatment of cancers is undoubted. Of equivalent importance is the theoretical work on clinical implications of cancer stem cells. Numerous models have shed light on clinical phenomena from a theoretical perspective and helped to explain patterns of treatment response and evolution of resistance [39, 46, 44, 30, 28].

Unfortunately, many models require involved parameterization which is implicitly difficult to obtain in a clinical setting. Their contribution to individualized treatment strategies thus remain unclear. In this work, we presented a very simple but general method to estimate the fraction of tumor driving cancer stem cells. This estimate can be made exclusively from the shape of a tumor’s growth curve. It only consists of a single exponential fit (linear regression) in the case of tumor growth, and two such regressions of longitudinal tumor size data during treatment.

The idea to estimate treatment prognosis and treatment response from biphasic tumor growth in leukemias is not novel. Different methods were suggested, yet they focus on the slope of the growth curve [41, 47]. Here, we show instead that the offset of the cancer growth regression, not the slope, allows for estimation of cancer stem cell fractions. Furthermore, we do not provide a method to estimate model parameters by fitting procedures, but show a direct functional link between two tumor properties, namely tumor growth and growth driving fraction of cancer stem cells.

Our method is parameter free. It requires no knowledge about microscopic properties of the tumor. It only utilizes techniques, for example high resolution images, which are already used routinely in clinical care. Thus, our method could readily complement current treatment protocols and inform about the relative size of the active pool of cancer stem cells.

Here, we neglect the potential emergence of treatment resistant sub-populations. However, the risk of the evolution of resistance depends critically on the size of the cancer driving stem cell pool size and the pre-existence of treatment resistant cells is much more likely than their spontaneous emergence during treatment [48]. Thus our method provides a tool to estimate the risk of a pre-existing treatment resistant sub population and might help to adjust treatment accordingly, for example a different combination of drugs.

Further, our model neglects a spatial component of tumor growth. This assumption leads to exponential growth in equilibrium, a situation well met in most leukemias [49, 50, 28], but also found in some solid tumors [51]. However, in some cases, the spatial component might be of importance and tumor growth becomes polynomial rather then exponential and our method provides only an approximation of the actual stem cell fraction. Yet, this divergence might be small compared to unavoidable errors induced by measurement related noise.

The ability to infer cancer stem cell fractions at diagnosis and during treatment can influence treatment strategies. Aggressiveness and duration of treatment might critically depend on the number of tumor driving cancer stem cells. In addition the risk of the evolution of resistance increases dramatically with an increasing stem cell pool [44]. Furthermore the composition of drug cocktails, as well as timing and scheduling could be adjusted according to knowledge gained about the tumor stem cell population. This potentially allows an opportunity to move away from the paradigm of maximum tolerated dose. Instead, it would provide a rational method to adaptively and a maximal effective dose for individualized treatment.

## 4 Methods

Here we lay out the deterministic dynamics of a hierarchically organized population. All cell divisions are symmetric. Stem cells divide at a rate *χ*_S_ and either self renew (producing two stem cells) with probability *p* or differentiate (producing two differentiated cells) with probability 1 – *p*. In addition stem cells might die at a rate *d*. Differentiated cells proliferate at a rate *χ*_D_ and die at a rate *d*. Further, differentiated cells can undergo a maximum of *m* cell doublings before they enter cell senescence. The system takes the form of a hierarchically coupled set of ordinary differential equations. The stem cell population obeys

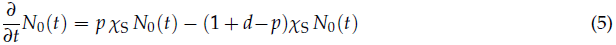

and the first compartment follows

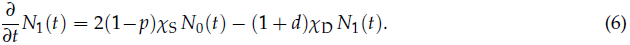

The function *N*_1_ corresponds to the number of differentiated cells that have not undergone further proliferations and thus have *m* cell cycles left before they enter cell cycle arrest. For all higher compartments (2 ≤ *i* ≤ *m*) we then have

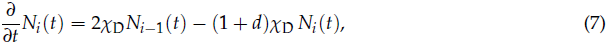

where *N*_*m*_ is the number of differentiated cells that only have a single cell cycle left after which they enter cell cycle arrest and are removed.

The equations (5)–(7) can be solved recursively for general initial conditions. If we set *N*_*i*_(0) to be the initial number of cells at time *t* = 0 in compartment *i*, we find

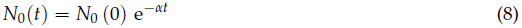

for the time dependence of cancer stem cells and *α* = (1 + *d* – 2*p*)*χ*_S_ is the net growth of cancer stem cells. The higher compartments (*i* > 0) evolve according to

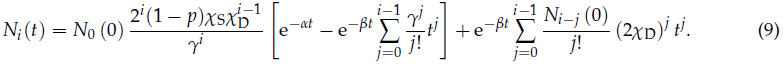

The outflow of each differentiated compartment is *β* = (1 + *d*)*χ*_D_ and *γ* = *β* – *α* corresponds to the differential outflow of stem and non stem cell compartments. The tumor transition from fast into slower growth is determined by the signs of *α* and *β* Both *χ*_D_ and *d* are strictly positive and consequently all terms in (9) that contain exp (–*βt*) vanish in the long run. If we have *p* > (1 + *d*)/2, *α* has a positive sign and determines tumor growth in the long run. Therefore, if we start from a single cell, *N*_0_(0) = 1, and the initial conditions become negligible, the total number of differentiated cells grows by

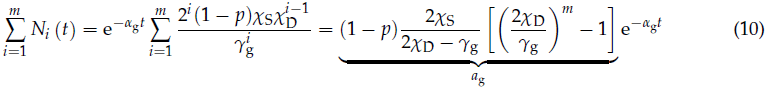

and follows a single exponential function *a*_g_e^−*α*_g_*t*^, with an offset *a*_g_ that involves all model parameters. The fraction of cancer stem cells can be written generally as

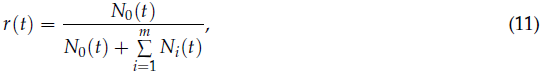

and is given in the slower growth phase by

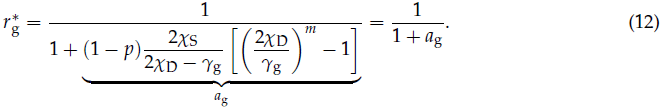

Similarly, during treatment, tumor growth in equilibrium can be written as N̄_t_ = *a*_t_e^−*α*_t_*t*^. However, in contrast to the tumor growth phase we get an additional term due to changed initial conditions. Instead of a single seeding cancer stem cell, we have e^*α*_g_*T*^ cancer stem cells at time of diagnosis *T*. Consequently, the fraction of stem cells becomes

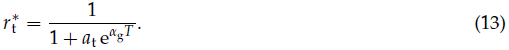

By re-substituting the age of the tumor via

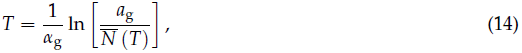

we find for the fraction of cancer stem cells under treatment

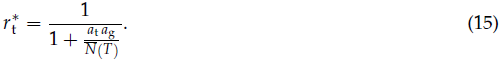

## 5 Acknowledgements

BW and AT acknowledge generous funding from the Max Planck Society. JGS would like to thank the NIH for its generous loan repayment grant. AS is supported by The Chris Rokos Fellowship in Evolution and Cancer. This work was also supported by the Wellcome Trust [105104/Z/14/Z]. PMA gratefully acknowledges financial support from Deutsche Akademie der Naturforscher Leopoldina, grant no. LPDS 2012-12. This paper was conceived during the 3rd annual Integrated Mathematical Oncology workshop on Personalized Medicine at the Moffitt Cancer Centre.

